# Prior expectations evoke stimulus templates in the deep layers of V1

**DOI:** 10.1101/2020.02.13.947622

**Authors:** Fraser Aitken, Georgios Menelaou, Oliver Warrington, Renée S. Koolschijn, Nadège Corbin, Martina F. Callaghan, Peter Kok

**Affiliations:** Wellcome Centre for Human Neuroimaging, UCL Queen Square Institute of Neurology, University College London, 12 Queen Square, London WC1N 3AR, UK; Wellcome Centre for Integrative Neuroimaging, University of Oxford, FMRIB, John Radcliffe Hospital, Oxford, OX3 9DU, UK

**Author notes:** **Corresponding author**, Wellcome Centre for Human Neuroimaging, UCL Queen Square Institute of Neurology, 12 Queen Square, London WC1N 3AR, UK.

**Keywords:** Perceptual inference, predictive coding, prediction, visual perception, layer fMRI, perceptual decision-making

## Abstract

The way we perceive the world is strongly influenced by our expectations. In line with this, much recent research has revealed that prior expectations strongly modulate sensory processing. However, the neural circuitry through which the brain integrates external sensory inputs with internal expectation signals remains unknown. In order to understand the computational architecture of the cortex, we need to investigate the way these signals flow through the cortical layers. This is crucial because the different cortical layers have distinct intra- and interregional connectivity patterns, and therefore determining which layers are involved in a cortical computation can inform us on the sources and targets of these signals. Here, we used ultra-high field (7T) functional magnetic resonance imaging (fMRI) to reveal that prior expectations evoke stimulus templates selectively in the deep layers of the primary visual cortex. These results shed light on the neural circuit underlying perceptual inference.

## Introduction

Much recent research has revealed that prior expectations strongly modulate sensory processing (Alink et al., 2010; De Lange et al., 2018; Den Ouden et al., 2009; Fiser et al., 2016; Gordon et al., 2019; Kok et al., 2012a; Summerfield et al., 2008; Todorovic et al., 2011; Yon et al., 2018), but it is as yet unclear what the neural mechanisms underlying these modulations are. To properly understand these mechanisms, we need to go beyond studying cortical regions as a whole, and study the laminar circuits involved in these computations (Lawrence et al., 2019b; Stephan et al., 2019). The reason for this is that the different cortical layers have different interregional connectivity patterns, with bottom-up signals predominantly flowing from superficial layers 2/3 to the granular layer 4 of downstream regions, and feedback arising from the deep layers 5/6 and targeting agranular layers 1 and 5/6 of upstream regions (Felleman and Van Essen, 1991; Harris and Mrsic-Flogel, 2013; Rockland and Pandya, 1979). Therefore, determining which layers are involved in a cortical computation can inform us on the likely sources and targets of these signals.

For instance, this known physiology has led to predictive coding theories (Bastos et al., 2012; Friston, 2005; Mumford, 1992; Rao and Ballard, 1999; Shipp, 2016) proposing that neurons in the deep cortical layers, which send feedback to upstream regions, represent our current hypotheses of the causes of our sensory inputs. Neurons in the superficial layers, on the other hand, are proposed to inform downstream regions of the mismatch between these hypotheses and current sensory inputs. While these theories are intriguing, and have garnered much excitement in the field (Clark, 2013; Keller and Mrsic-Flogel, 2018), there has been no direct empirical support of these different proposed roles of the cortical layers.

One recent proposal is that expectations evoke stimulus templates in the primary visual cortex (Hindy et al., 2016; Kok et al., 2014, 2017), which in turn modulate processing of subsequent sensory inputs (Summerfield and de Lange, 2014). The goal of the current study was to determine which layers of the primary visual cortex (V1) contain such expectation templates, and how this differs from the activity evoked by a stimulus presented to the eyes. We hypothesised that merely expecting a stimulus might activate a template of the expected stimulus in the deep layers of the visual cortex, which have been proposed to contain perceptual hypotheses (Friston, 2005; Kok et al., 2016; Lee and Mumford, 2003; Shipp, 2016). Alternatively, expectations may serve to increase the synaptic gain on expected sensory signals in superficial layers, akin to mechanisms of feature-based attention (Lawrence et al., 2019a; Martino et al., 2015).

## Results

We induced expectations by presenting a cue (orange vs. cyan circle inside a fixation bull’s eye) that predicted the orientation (45° vs. 135°) of a subsequently presented grating stimulus (Fig. 1a). This first grating was followed by a second grating which differed slightly in orientation (mean = 3.3°) and contrast (mean = 6.9%), determined by an adaptive staircase (see Experimental Procedures). In separate runs of the experiment, human participants (N=18) performed two tasks, judging either the orientation or contrast change between the two gratings. Crucially, on 25% of trials, the gratings were omitted (Fig. 1b), meaning the screen stayed empty except for the fixation bull’s eye, and participants were not required to perform any task. Therefore, on these trials, participants had a highly specific expectation of a certain stimulus appearing, but there was no corresponding input to the eyes.

**Figure 1.**
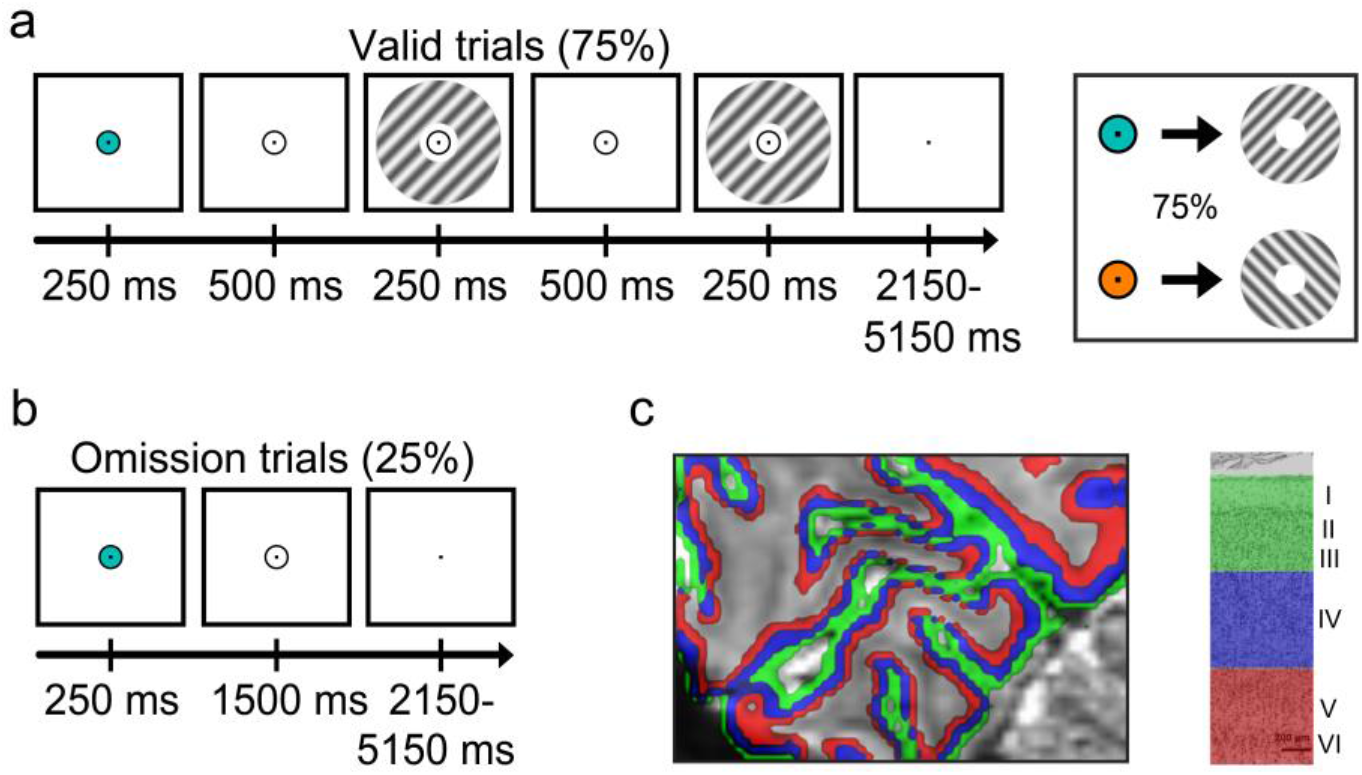
Experimental paradigm. **a,** Each trial started with a coloured dot (cyan or orange) that predicted the orientation of the subsequent grating stimulus (45° or 135°). On 75% of trials a set of gratings was then presented, the first of which had the (expected) orientation and the second differed slightly in orientation and contrast. In separate experimental runs, participants discriminated either the orientation or the contrast difference between the gratings. **b,** In 25% of trials, the gratings were omitted. On these trials, there was an expectation of a particular visual stimulus but no actual visual input. Participants had no task in these trials, except for holding central fixation. **c,** Sagittal slice of the mean functional scan of the occipital lobe of one participant. Deep, middle and superficial cortical layers indicated in coloured ribbons. Cytoarchitectural image of V1 adapted from (de Sousa et al., 2010).

We non-invasively examined the laminar profile of the activity evoked by these orientation expectations in human V1, using ultra-high-field (7T) fMRI with high spatial resolution (0.8 mm isotropic). Layer-specific fMRI is a novel technique that has only recently become feasible due to the sub-millimetre resolution required. It has been successfully used to study many cognitive processes, such as attention (Klein et al., 2018; Lawrence et al., 2019a; Martino et al., 2015), working memory (Finn et al., 2019; Lawrence et al., 2018), spatial context (Muckli et al., 2015), perceptual illusions (Kok et al., 2016), and even language (Sharoh et al., 2019). Encouragingly, results have generally been in good alignment with those obtained from invasive animal studies (Fracasso et al., 2016; Huber et al., 2017; Self et al., 2019).

To examine orientation-specific blood-oxygen-level dependent (BOLD) activity, we divided V1 voxels into two (45°-preferring and 135°-preferring) sub-populations depending on their orientation preference during an independent functional localiser. Layer-specific time courses were extracted for both voxel sub-populations. Specifically, we defined three equivolume gray matter layers (superficial, middle, and deep; Fig. 1c) and determined the proportion of each voxel’s volume in these layers (as well as in white matter and cerebrospinal fluid). These layer “weights” were subsequently used in a spatial regression analysis to estimate layer-specific time courses of the BOLD signal in the two sub-populations of V1 voxels (Kok et al., 2016; Lawrence et al., 2019a, 2018; Van Mourik et al., 2019). This regression analysis served to unmix the signals originating from the different layers, which were potentially mixed within individual voxels. Finally, for both voxel sub-populations, we subtracted the activity of the three layers evoked by expecting/seeing the non-preferred orientation from the activity evoked by expecting/seeing the preferred orientation. This procedure resulted in orientation-specific, layer-specific BOLD signals (Lawrence et al., 2019a, 2018) for both the trials on which gratings were presented, and those on which the gratings were expected but omitted.

### Prior expectations selectively activate deep layers of V1

The laminar profile of V1 activity evoked by purely top-down expectations, in the absence of bottom-up input, was strikingly different from that evoked by actually presented stimuli (Fig. 2; interaction between presented vs. omitted and cortical layer: *F*_2,34_ = 5.4, *p* = 0.0093). Specifically, merely expecting a grating with a specific orientation evoked a BOLD signal reflecting that orientation in the deep (*t*_17_ = 3.5, *p* = 0.0029), but not the middle (*t*_17_ = 0.8, *p* = 0.45) or superficial (*t*_17_ = 0.7, *p* = 0.48) layers of V1. When a grating was actually presented to the eyes, this evoked orientation-specific activity in all layers of V1 (deep: *t*_17_ = 4.5, *p* = 0.00035; middle: *t*_17_ = 3.6, *p* = 0.0022; superficial: *t*_17_ = 4.2, *p* = 0.00063), as would be expected.

**Figure 2.**
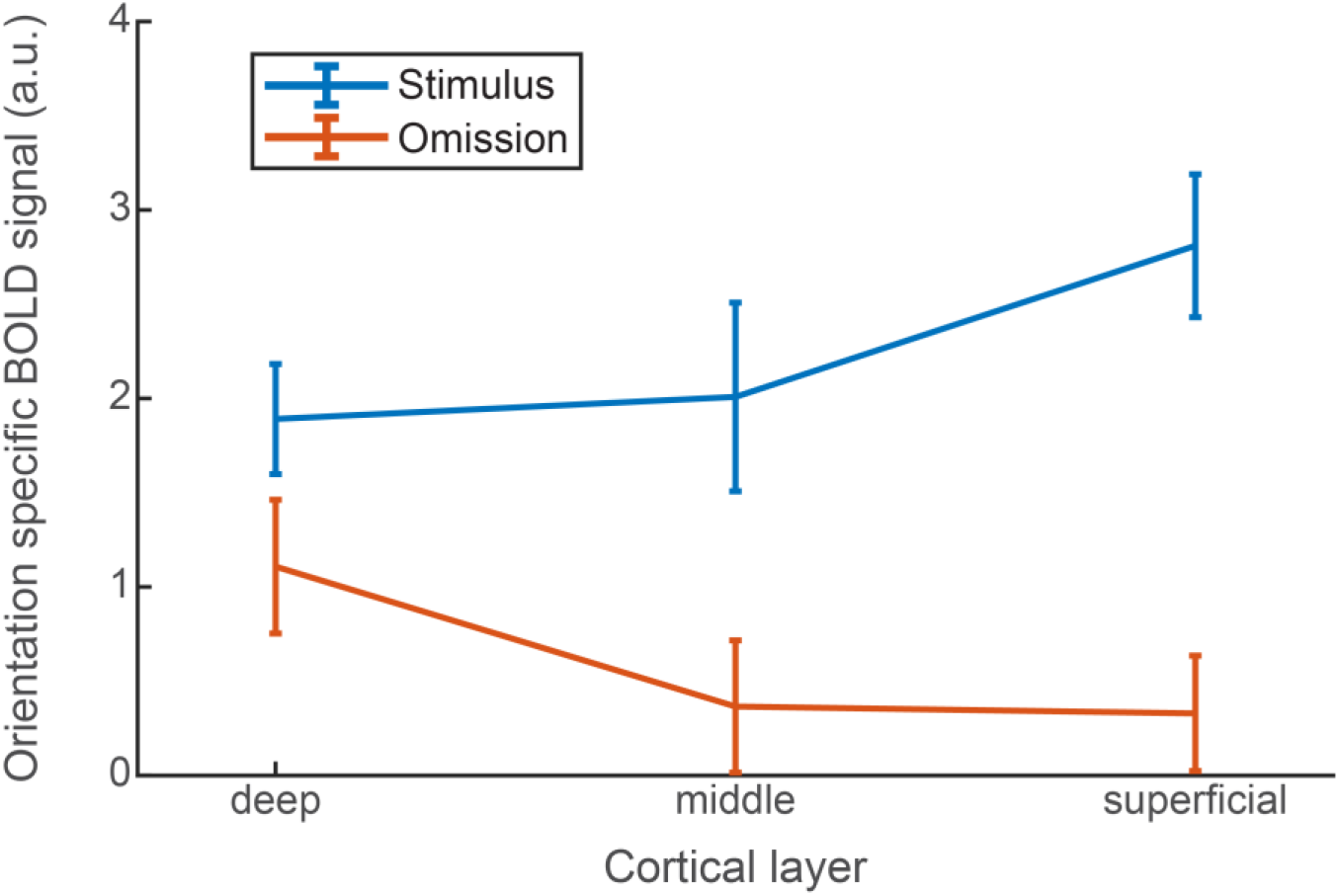
Layer-specific BOLD response in V1 for presented and expected stimuli. Orientation-specific BOLD response to presented (blue) and expected-but-omitted (orange) gratings in the different layers of V1. Error bars indicate within-subject SEM.

Interestingly, the strength of the expectation signals in the omission trials was dependent on the task participants were performing in that experimental run (Fig. 3; interaction between task and cortical layer: *F*_2,34_ = 3.6, *p* = 0.039). That is, orientation expectations evoked a stimulus template in the deep layers of V1 when participants were preparing to perform an orientation discrimination (*t*_17_ = 3.0, *p* = 0.0086), but not when they were preparing to discriminate the contrast of the gratings (*t*_17_ < 0.1, *p* = 0.98; though note that there was no significant difference between tasks in the deep layers: *t*_17_ = 1.5, *p* = 0.16). Given that accuracy and reaction times did not differ between the two tasks (accuracy: 79.6% vs. 78.9%; *t*_17_ = 0.4, *p* = 0.72; RT: 689 ms vs. 676 ms; *t*_17_ = 1.2, *p* = 0.23), this is unlikely to be due to task difficulty or engagement, but more likely due to the fact that the expected feature, orientation, was task-relevant in one task but not the other. We are cautious to over interpret this effect given that some previous studies with similar experimental designs have not shown it (Kok et al., 2012a, 2017), but this could indicate that expectation signals are stronger when they pertain to a task-relevant or attended feature (Richter and De Lange, 2019).

**Figure 3.**
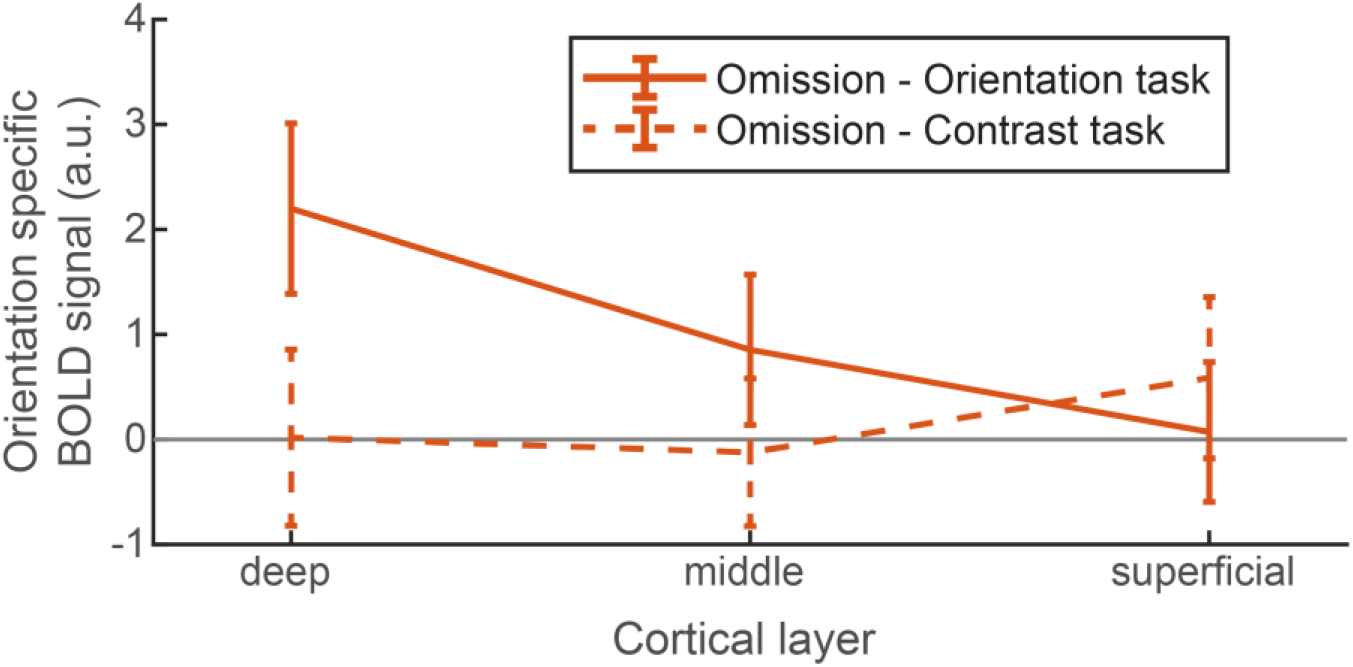
Layer-specific BOLD response in V1 for expected stimuli, per task. Orientation-specific BOLD response to expected-but-omitted gratings in the different layers of V1, separately for experimental runs in which participants performed the orientation (solid lines) and contrast (dashed lines) discrimination tasks. Error bars indicate within-subject SEM.

## Discussion

In short, we found that prior expectations evoke stimulus-specific activity selectively in the deep layers of V1. Interestingly, this selective activation of the deep layers matches that evoked by illusory Kanizsa figures, which have been suggested to be the result of an automatic structural expectation in the visual system (Kok et al., 2016). However, since the expectations in the current experiment were signalled by a conditional cue, V1 activity is more likely to be the result of feedback from higher-order regions outside of the visual cortex, possibly involving the hippocampus (Hindy et al., 2016; Kok and Turk-Browne, 2018; Kok et al., 2020; Schapiro et al., 2012). The fact that these very different types of ‘expectations’ evoke highly similar layer-specific activity in visual cortex may point to a common computational role for these expectations in V1.

Specifically, one proposed implementation of perceptual inference is predictive coding (Friston, 2005; Rao and Ballard, 1999), a theory that proposes that each cortical region houses separate sub-populations of neurons coding for perceptual hypotheses (predictions) and mismatches between these hypotheses and bottom-up sensory input (prediction errors). Since feedback mainly arises from the deep layers, prediction units are suggested to reside predominantly in the deep layers, while prediction error units dominate in the middle and superficial layers (Bastos et al., 2012; Shipp, 2016). The current results are in line with this proposed arrangement, since a prediction in the absence of any bottom-up input was found to evoke stimulus-specific signals in only the deep layers of V1.

Notably, V1 templates evoked by maintaining a grating stimulus in working memory have a strikingly different laminar profile, activating both the deep and the superficial (but not the middle) layers (Lawrence et al., 2018). The recruitment of the deep layers by both expectation and working memory could represent the use of internally generated stimulus templates in both processes. In addition, the conscious effort of maintaining a stimulus in working memory for goal-directed behaviour may require feature-based attention, which has been suggested to modulate activity in the superficial layers (Lawrence et al., 2019a; Martino et al., 2015). Therefore, working memory may be a compound process, activating deep and superficial layers for different computational reasons. More broadly, it has been suggested that expectation and attention may be separable neural processes (Gordon et al., 2019; Summerfield and Egner, 2009, 2016), with distinct computational roles: activating hypotheses and boosting relevant inputs, respectively (Feldman and Friston, 2010; Kok et al., 2012b). Layer-specific fMRI offers the opportunity to directly address these questions in the human brain for the first time (Stephan et al., 2019).

It is interesting to note that the presence of the expectation templates depended on the task participants were performing, suggesting that expectation signals were stronger when they pertained to an attended feature (i.e., during the orientation task) than to an unattended feature (during the contrast task) (Richter and De Lange, 2019). (Though note that the deep layer templates were not significantly stronger in the orientation task than in the contrast task, prompting caution in interpreting these effects.) This dependency was not found in recent studies using a very similar experimental design investigating expectation templates using magnetoencephalography (Kok et al., 2017) and effects of expectation on stimulus processing in V1 using fMRI (Kok et al., 2012a). One possibility is that these previous studies simply were not able to detect these task modulations because they lacked laminar resolution; note that the task dependence here is expressed as an interaction between task and cortical layer. An alternative possibility is that attention may boost the gain of expectation templates, perhaps even promoting them from activity-silent synaptic templates to being reflected in neural firing (Rose et al., 2016; Wolff et al., 2017), but that this does not change the consequences of these templates for subsequent stimulus processing (Kok et al., 2012a). Future layer-specific research orthogonally manipulating attention and expectation validity will be needed to distinguish these possibilities.

How might stimulus templates in the deep layers modulate processing of incoming sensory inputs? One potential mechanism for this is through inhibitory connections from the deep layers to the middle and superficial layers (Harris and Mrsic-Flogel, 2013; Kätzel et al., 2011; Thomson and Bannister, 2003), which might cause a reduction in activity throughout the entire cortical column (Olsen et al., 2012) as a result of the excitatory pathways from layer 4 to layers 2/3 and from there to layers 5/6 (Felleman and Van Essen, 1991; Harris and Mrsic-Flogel, 2013). In the absence of sensory input to layer 4, as is the case in the omission trials here, this modulation would not occur and top-down feedback signals would be restricted to the deep layers. An alternative mechanism could be through feedback connections terminating on inhibitory neurons in layer 1, which in turn inhibit pyramidal neurons in layers 2/3 (Bastos et al., 2012; Chu et al., 2003). Possibly in line with this, a recent laminar fMRI study of contextual effects in non-stimulated human V1 reported effects in the most superficial layers, potentially reflecting layer 1 (Muckli et al., 2015). It should be noted that layer 1 is difficult to detect with layer-specific fMRI, as it is very thin and sparsely populated with neurons.

The finding that prior expectations evoke stimulus templates selectively in the deep layers of visual cortex sheds light on the neural circuit by which the brain performs perceptual inference (Bastos et al., 2012; Lee and Mumford, 2003; Shipp, 2016); that is, combines sensory signals with internal expectations to generate a best guess of what is out there in the world. Ultimately, future work building on these findings may be able to reveal how this delicate balance between internal and external signals can go awry, as it does in disorders such as autism (Lawson et al., 2017; Utzerath et al., 2018; Van de Cruys et al., 2014) and psychosis (Corlett et al., 2019; Powers et al., 2017; Stephan et al., 2019).

## Experimental procedures

### Participants

23 healthy human volunteers with normal or corrected-to-normal vision participated in the 7T fMRI experiment. The study was approved by the Oxford University Ethics Committee, and all participants gave informed consent and received monetary compensation. One participant was excluded because they responded to < 50% of trials during the fMRI session. One further participant was excluded because the calcarine sulcus was not in the field of view for the entire fMRI session, due to large head movements between runs. Finally, three participants were excluded due to our strict head motion criteria of no more than 10 movements larger than 1.0 mm between successive functional volumes. The final sample consisted of 18 participants (10 female; age 25 ± 4 years; mean ± SD).

### Stimuli

Grayscale luminance-defined sinusoidal Gabor grating stimuli were generated using MATLAB (MathWorks, Natick, MA, USA, RRID:SCR_001622) and the Psychophysics Toolbox (Brainard, 1997). During the behavioural session, the stimuli were presented on an Apple MacBook Pro (1280 x 800 screen resolution, 60 Hz refresh rate). In the fMRI scanning session, stimuli were projected onto a rear projection screen using an Eiki LC-XL100 projector with custom throw lens (1024 x 768 screen resolution, 60 Hz refresh rate) and viewed via a mirror (view distance 60 cm). Visual prediction cues consisted of a circular region within a white fixation bull’s eye (0.7° diameter) turning either cyan or orange for 250 ms. On valid trials (75%), cues were followed by a set of two gratings (1.5 cpd spatial frequency, 250 ms duration each, separated by a 500 ms blank screen), displayed in succession in an annulus (outer diameter: 10° of visual angle, inner diameter: 1°, contrast decreasing linearly to zero over 0.7° at the inner and outer edges), surrounding a fixation bull’s eye (0.7° diameter). The central fixation bull’s eye was presented throughout the trial, as well as during the inter-trial interval (ITI, jittered exponentially between 2150 and 5150 ms).

### Experimental procedure

Trials consisted of a coloured prediction cue, followed by two consecutive grating stimuli on 75% of trials (750 ms stimulus onset asynchrony (SOA) between cue and first grating) (Fig. 1a). The coloured cue (cyan or orange) predicted the orientation of the first grating stimulus (45° or 135°) (Fig. 1a). On valid trials (75%), two consecutive grating stimuli were presented following the coloured cue. The first grating had the orientation predicted by the cue (45° or 135°), and a luminance contrast of 80%. The second grating differed slightly from the first in terms of both orientation and contrast (see below), as well as being in antiphase to the first grating (which had a random spatial phase). On the remaining 25% of trials, no gratings were presented (omission trials; Fig. 1b), and the screen remained empty except for the fixation bull’s eye. Participants had no task on these trials, except for holding central fixation. The contingencies between the cue colours and grating orientations were flipped halfway through the experiment (i.e., after two runs), and the order was counterbalanced over participants.

In separate runs (two blocks of 64 trials each, ~13 minutes), participants performed either an orientation or a contrast discrimination task on the two gratings. When performing the orientation task, participants had to judge whether the second grating was rotated clockwise or anticlockwise with respect to the first grating. In the contrast task, a judgment had to be made on whether the second grating had lower or higher contrast than the first one. These tasks were explicitly designed to avoid a direct relationship between the perceptual expectation and the task response. Furthermore, these two different tasks were designed to manipulate the task-relevance of the grating orientations, to investigate whether the effects of orientation expectations depend on the task-relevance of the expected feature. Participants indicated their response (response deadline: 750 ms after offset of the second grating) using an MR-compatible button box. The orientation and contrast differences between the two gratings were determined by an adaptive staircase procedure(Watson and Pelli, 1983), being updated after each trial. This was done to yield comparable task difficulty and performance (~ 75% correct) for the different tasks. Staircase thresholds obtained during one task were used to set the stimulus differences during the other task, in order to make the stimuli as similar as possible in both contexts.

All participants completed four runs (two of each task, alternating every run, order was counterbalanced over participants) of the experiment, yielding a total of 512 trials, 256 per task. At the start of each block, the relationship between the cue and the stimulus was shown by presenting the predicted orientation within an appropriately coloured circle. The staircases were kept running throughout the experiment. Prior to entering the scanner, as well as in between runs two and three, when the contingencies between cue and stimuli were flipped, participants performed a short practice run containing 32 trials of both tasks (~4.5 minutes).

Participants underwent a behavioural practice session just prior to entering the scanner to ensure knowledge of the task and how to respond. In the practice session, participants were given written and verbal instructions about the task requirements. During the practice runs, the coloured cues predicted the orientation of the first grating stimulus of the pair with 100% validity (45° or 135°; no omission trials).

After the main experiment, participants performed a functional localizer task inside the scanner. This consisted of flickering gratings (2 Hz), presented at 100% contrast, in blocks of ~14.3 seconds (4 TRs). Each block contained gratings with a fixed orientation (45° or 135°). The two orientations were presented in a pseudorandom order followed by a ~14.3 s blank screen, containing only a fixation bull’s eye. Participants were tasked with responding whenever the white fixation dot briefly dimmed, to ensure central fixation. All participants were presented with 16 localiser blocks.

### fMRI data acquisition

Functional images were acquired on a Siemens Magnetom 7T MRI system (Siemens Healthcare GmbH, Erlangen, Germany) with a single channel head coil for localised transmission with a 32-channel head coil insert for reception (Nova Medical, Wilmington, USA) at the Wellcome Centre for Integrative Neuroimaging Centre (University of Oxford), using a T2*-weighted 3D gradient-echo EPI sequence (volume acquisition time of 3583 ms, TR = 74.65 ms, TE = 29.75 ms, voxel size 0.8×0.8×0.8 mm, 16° flip angle, field of view 192 × 192 × 38.4 mm, in-plane GRAPPA acceleration factor 4, in-plane partial Fourier 6/8). Anatomical images were acquired using an MPRAGE sequence (TR = 2200 ms, TE = 2.96 ms, TI = 1050 ms, voxel size 0.7×0.7×0.7 mm, 7° flip angle, field of view 224 × 224 × 179.2 mm, in-plane GRAPPA acceleration factor 2).

### Preprocessing of fMRI data

The first four volumes of each run were discarded. The functional volumes were cropped to cover only the occipital lobe to reduce the influence of severe distortions in the frontal lobe. The cropped functional volumes were spatially realigned within scanner runs, and subsequently between runs, to correct for head movement using SPM12. FSL FAST (Zhang et al., 2001) was used to correct the bias field and remove intensity gradients in the MPRAGE anatomical image.

### Segmentation and coregistration of cortical surfaces

Freesurfer (http://surfer.nmr.mgh.harvard.edu/) was used to detect boundaries between grey and white matter and cerebrospinal fluid (CSF), respectively, on the basis of the bias-corrected MPRAGE. Manual corrections were made to remove dura incorrectly included in the pial surface when necessary. The gray matter boundaries were registered to the mean functional volume in two steps: 1) a conventional rigid body boundary-based registration (BBR) (Greve and Fischl, 2009), and 2) recursive boundary registration (RBR) (Mourik et al., 2019). During RBR, BBR was applied recursively to increasingly smaller partitions of the cortical mesh. Here, we applied affine BBR with 7 degrees of freedom: rotation and translation along all three dimensions, and scaling along the phase-encoding direction. In each iteration, the cortical mesh is split into two, the optimal BBR transformations are found and applied to the respective parts. Subsequently, each part is split into two again and registered. The specificity increases at each stage, and corrects for local mismatches between the structural and the functional volumes that are due to magnetic field inhomogeneity related distortions. Here, we ran six such iterations. The splits are made along the cardinal axes of the volume, such that the number of vertices is equal for both parts. The plane for the second cut is orthogonal to the first, the third orthogonal to the first two. The median displacement was taken after running the recursive algorithm six times, in which different splitting orders where used, comprised of all six permutations of x, y and z.

### Definition of regions of interest

The V1 surface label, defined by Freesurfer based on the anatomy of the MPRAGE image, was projected into volume space covering the full cortical depth plus a 50% extension into WM and CSF, respectively. The V1 region of interest (ROI) was constrained to only the most active voxels in response to the grating stimuli by applying a temporal General Linear Model (GLM) to the preprocessed data from the functional localiser run. Blocks of 45° and 135° gratings were modelled separately as regressors and contrasted against baseline to identify voxels that exhibited a significant response to the grating stimuli (*t* > 2.3, *p* < 0.05). To estimate the orientation preference of each voxel, the two orientation regressors were contrasted against each other. We selected the 500 voxels with the most positive T values (45° preferring) and the 500 voxels with the most negative T values (135° preferring) in the contrast, and created separate masks for each. Finally, we z-scored the time course data from each voxel and multiplied this time course with the absolute T-value from the orientation contrast (45° vs. 135°) in order to weight the results towards the voxels with the most robust orientation preference (Lawrence et al., 2019a, 2018).

### Definition of the cortical layers

Gray matter was divided into three equivolume layers using the level-set method (described in detail in (Kleinnijenhuis et al., 2015; Van Mourik et al., 2019; Waehnert et al., 2014)) following the principle that the layers of the cortex maintain their volume ratio throughout the curves of the gyri and sulci (Bok, 1929). We calculated four level set functions creating five distinct cortical compartments: white matter, three gray matter layers (deep, middle and superficial) and CSF. In human V1, these three laminar compartments are expected to correspond roughly to layers I–III, layer IV, and layers V–VI, respectively (de Sousa et al., 2010) (Fig. 1c). The level set functions allowed calculation of a laminar design-matrix describing the distribution of each voxel’s volume in an ROI over the five compartments (Kok et al., 2016; Lawrence et al., 2019a, 2018; Van Mourik et al., 2019).

### Extraction of layer-specific time courses

To reduce partial-volume effects and improve the spatial resolution, the laminar design-matrix can be used in a spatial GLM to separate the voxels’ BOLD signal for each of the five compartments (Van Mourik et al., 2019):

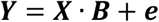

**Y** is a vector of voxel values from an ROI, **X** is the laminar design-matrix and **B** is a vector of layer signals. For each ROI and each functional volume, the layer signal **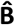** was estimated by regressing **Y** against **X**, yielding five depth-specific time courses per ROI and functional run.

### Estimating effects of interest per layer

We estimated the effects of interest in the three gray matter layers using a temporal GLM. Regressors of interest (presentation/omission of 45° or 135° gratings, separately for orientation and contrast task runs) were constructed by convolving stick functions representing the onsets of the trials with SPM12’s canonical haemodynamic response function. Regressors of no interest included head motion parameters, their derivatives and the square of the derivatives. Both the data and design matrix were high-pass filtered (cutoff = 128 s) to remove low-frequency signal drifts.

To calculate orientation-specific BOLD responses, for each ROI (e.g., the 45°-preferring V1 ROI) the estimated BOLD response for conditions in which the non-preferred orientation was presented/expected (e.g., a 135° grating expected-but-omitted) was subtracted from the response for the corresponding condition in which the preferred orientation was presented/expected (e.g., a 45° grating expected-but-omitted). After this subtraction, responses were averaged over the two V1 ROIs, yielding layer-specific orientation-specific BOLD responses to each of the conditions of interest (expected-and-presented and expected-but-omitted gratings, per task). Our main effect of interest, namely whether laminar BOLD profiles differed for presented and expected-but-omitted stimuli, was tested using 2×3 repeated-measures ANOVA with factors stimulus type (presented vs. omitted) and cortical layer (deep, middle and superficial). To investigate whether expectation effects were task-dependent, a 2×3 repeated-measures ANOVA with factors task (orientation vs. contrast task) and cortical layer (deep, middle and superficial) was conducted. Significant interactions were followed up with paired-sample t-tests.

## Acknowledgements

The authors would like to thank the Wellcome Centre for Integrative Neuroimaging radiography team for assistance with data collection, and Nancy Rawlings for coordinating project administration. We are grateful to Stephen Fleming and Floris de Lange for comments on an earlier version of the manuscript. This work was supported by a Sir Henry Dale Fellowship jointly funded by the Wellcome Trust and the Royal Society (218535/Z/19/Z) to P.K. R.S.K. is supported by an EPSRC/MRC-funded studentship (EP/L016052/1). The Wellcome Centre for Human Neuroimaging is supported by core funding from the Wellcome Trust (203147/Z/16/Z). The Wellcome Centre for Integrative Neuroimaging is supported by core funding from the Wellcome Trust (203139/Z/16/Z).

## Author contributions

P.K. designed the study; N.C. and M.F.C. developed the fMRI scanning sequence; F.A., G.M., and R.S.K. collected the data; F.A., O.W., and P.K. analysed the data; all authors wrote the manuscript.

